# Co-occurrence of Antibiotic Residues and Antimicrobial Resistance Genes in Animal Manure and Agricultural Soils from Machakos, Kiambu, and Kajiado Counties, Kenya

**DOI:** 10.1101/2025.11.09.687469

**Authors:** Richard M. Machoka, Florence Ng’ong’a, Christabel Muhonja, Heshborne Tindih, Zachary Getenga, Thiele-Bruhn Sören

## Abstract

The excessive and often unregulated use of antibiotics in livestock production and human health has led to the dissemination of antibiotic residues and antibiotic resistance genes (ARGs) posing a serious threat to the environment and public health. This study investigates the co-occurrence and spatial distribution of antibiotic residues and ARGs in livestock manure and agricultural soils from Machakos, Kiambu, and Kajiado counties in Kenya, regions characterized by intensive livestock farming. A total of 180 samples from 30 farms across the three counties were collected and pooled into 18 samples. Antibiotic residues were extracted and quantified using high-performance liquid chromatography coupled with tandem mass spectrometry (LC-MS/MS). Nine ARGs (*aadA, ermB*, *sul1, tetQ, tetW, dfrA1, blaMOX, blaOXA* and *qnrB* were quantified via absolute and relative qPCR, normalized to *16s rRNA* gene copy numbers. Multivariate analyses, including PCoA and Spearman Correlation, were conducted to explore ARG structure and co-occurrence. Tetracycline, particularly oxytetracycline, was the most abundant antibiotic (up to 1150 ng/mL), especially in pig manure from Kiambu. Sulfadimethoxin was undetectable in nearly all samples except in soil mixed with pig manure from Kajiado county, which showed a concentration above 70 ng/mL, the pig manure from the same county had levels below 2 ng/mL. Sulfadimidin, sulfamethoxanol and erythromycin were not detected in all samples. A three-way ANOVA showed that animal source and sample type were generally not significant factors in antibiotic concentration, but oxytetracycline (p= 0.0421, 0.0901) and sulfadimethoxin (P =0.0904, 0.044, 0.054) showed marginal significance by sample type (P=0.0785). Antibiotic resistance genes were widely distributed, *with aadA, ermB* and *sul1* being the most prevalent, especially in soils from Kiambu and Machakos. *TetQ* showed extremely high relative abundance, indicating intense tetracycline selection pressure. Strong positive correlations were observed between co-occurring ARGs, including *tetQ* and *tetW and erm* and *sul1*. The concurrent detection of persistent antibiotic residues and high ARG loads in both manure and soils underscores the urgent need for improved antibiotic stewardship, sustainable manure management, and environmental monitoring in Kenyan agroecosystems.

## Background

Antibiotics are commonly used in the treatment of animal and human diseases; however, excessive usage of antibiotics has led to an increase in antibiotic-resistant microbes and antibiotic-resistant genes (ARGs), which has become a threat to human and animal health (Dutta, 2025). Thus, the World Health Organization (WHO) has listed antimicrobial resistance among the top ten public health threats facing humanity (Tejpar et al., 2022). Globally, about 700,000 deaths result from infections caused by antimicrobial-resistant pathogens annually, with more deaths occurring in Sub-Saharan Africa (Kariuki et al., 2022). This number may rise to about 10 million by 2050 if nothing is done urgently, (Murray et al., 2022; Shankar, 2016).

In veterinary medicine, antibiotics consists of over 70% of all pharmaceuticals in livestock production (Thiele-Bruhn, 2003) leading to the spread of antimicrobial resistance, especially when used as prophylactic drugs (Checcucci et al., 2020). The antibiotics used in low doses in animals to promote growth provide a perfect growth environment for antimicrobial-resistant pathogens (Chantziaras et al., 2014; Pokharel et al., 2020). In this regard, animals may act as a reservoir for antibiotic-resistant genes (ARGs), which then may be transferred to the environment via excrement and consequently to humans through direct handling of animals and or the food chain (Gao et al., 2020; Pokharel et al., 2020). The cycle of consumption and contamination is facilitated through pollution, organic fertilizers, industrial waste, and poor sanitation, contributing to the environmental resistome. The role and significance of animal waste in AMR spread in the environment is therefore crucial.

Livestock excrete between 40 - 90% of the antibiotics either as active metabolites or unchanged isomers or epimers of the parent molecule in feces and urine (Polianciuc et al., 2020). Some metabolites end up being more potent than the parent molecule, while others can change back to their parent compounds during manure storage, for example, sulphonamides and acetic conjugates (Massé et al., 2014) The concentration of these antibiotics in manure vary between 1-10mgkg^-1^ or L^-1,^ although concentrations as high as 200 mgkg^-1^ or L^-1^ have been reported (Kumar et al., 2005).

In recognition of the public health importance of AMR to both human and the environment and in alignment with the Global Action Plan on AMR, the Kenyan government released a National Action Plan on AMR in 2017, which was revised in 2023 to formulate a comprehensive response to AMR by involving all relevant stakeholders (Gitonga et al., 2024). In this report, it is recognized that combating AMR requires a One Health approach from all sectors to address the gaps in the fight against AMR, which include a lack of knowledge, poor surveillance, and misuse of antibiotics by farmers (Mukoko et al., 2025).

About 90% of all antibiotic purchases in Kenya for veterinary purposes are used for prophylactic activities rather than curative (WHO, 2022). The Kenya Veterinary Board estimated that over 33% of antibiotics available for animal use are substandard or counterfeit, complicating the situation further (Sohaili et al., 2024). Antibiotic residues have been found in various animal products sold in markets in Kenya, including chicken meat (Odundo et al., 2023), milk sold by vendors in Kibera, where beta-lactams and oxytetracyclines were found in over 66% of unpasteurized milk (Brown et al., 2020), Sulfonamide residues have been isolated from the dairy value chain in Nakuru, Kenya (Orwa et al., 2017).

Therefore, there is a need to understand the concentrations of various antibiotic residues and the ARG profiles from different animal manure and agricultural soils in Kenya. This study focused on 3 counties in Kenya that are highly agricultural with a routine usage of antibiotics for various animals.

## Materials and methods

### Study site

This study was conducted in three counties in Kenya: Machakos, Kiambu and Kajiado. Machakos borders Nairobi and Kiambu Counties to the west, Kitui County to the East, Makueni county to the South, Embu County to the North Kajiado to the southwest. It is located in latitude 0° 45’ South to 1°31’ South and longitudes 36° 45’ East to 37° 45’ East. Administratively, Machakos is divided into about 40 wards and it covers about 6,208 KM^2^ with an estimated population of 1,421,932. Subsistence farming is the main economic activity in the county where mixed farming is practiced (Kenya National Bureau of Statistics, 2015b).

Kiambu County is located in the central region and covers a total area of 2,543.5 Km^2^ with 476.3 Km^2^ under forest cover according to the 2009 Kenya Population and Housing Census. Kiambu County borders Nairobi and Kajiado Counties to the South, Machakos to the East, Murang’a to the North and northeast, Nyandarua to the northwest, and Nakuru to the West. The county lies between latitudes 00 25’and 10 20’South of the Equator and Longitude 360 31‘and 370 15‘East. It has a population of about 1,760,692 people. Small-scale farming is the main economic activity and zero grazing is practiced given the limited farm sizes in the county. Animals reared include cattle, Sheep and goats, poultry and pigs. Growth in livestock farming has been motivated by the ready market in the urban centres and the availability of local food processing factories(Statistics, 2015).

Kajiado County covers an approximate area of 21,900.9 square kilometers (Km^2^). It is located in the southern part of Kenya and borders Nairobi County to the North East, Narok County to the West, Nakuru and Kiambu Counties to the North, Taita Taveta County to the South East, Machakos and Makueni Counties to the North East and East respectively, and the Republic of Tanzania to the South. It is situated between Longitudes 360 5’ and 370 5’ East and between Latitudes 10 0’ and 30 0’ South. The main economic activities carried out mainly in Kajiado are pastoralist, wholesale and retail trade, mining, especially soda ash in Magadi and marble in Kajiado central, and agriculture, which includes both horticulture and small-scale farming(Kenya National Bureau of Statistics, 2015a).

**Figure 1:**
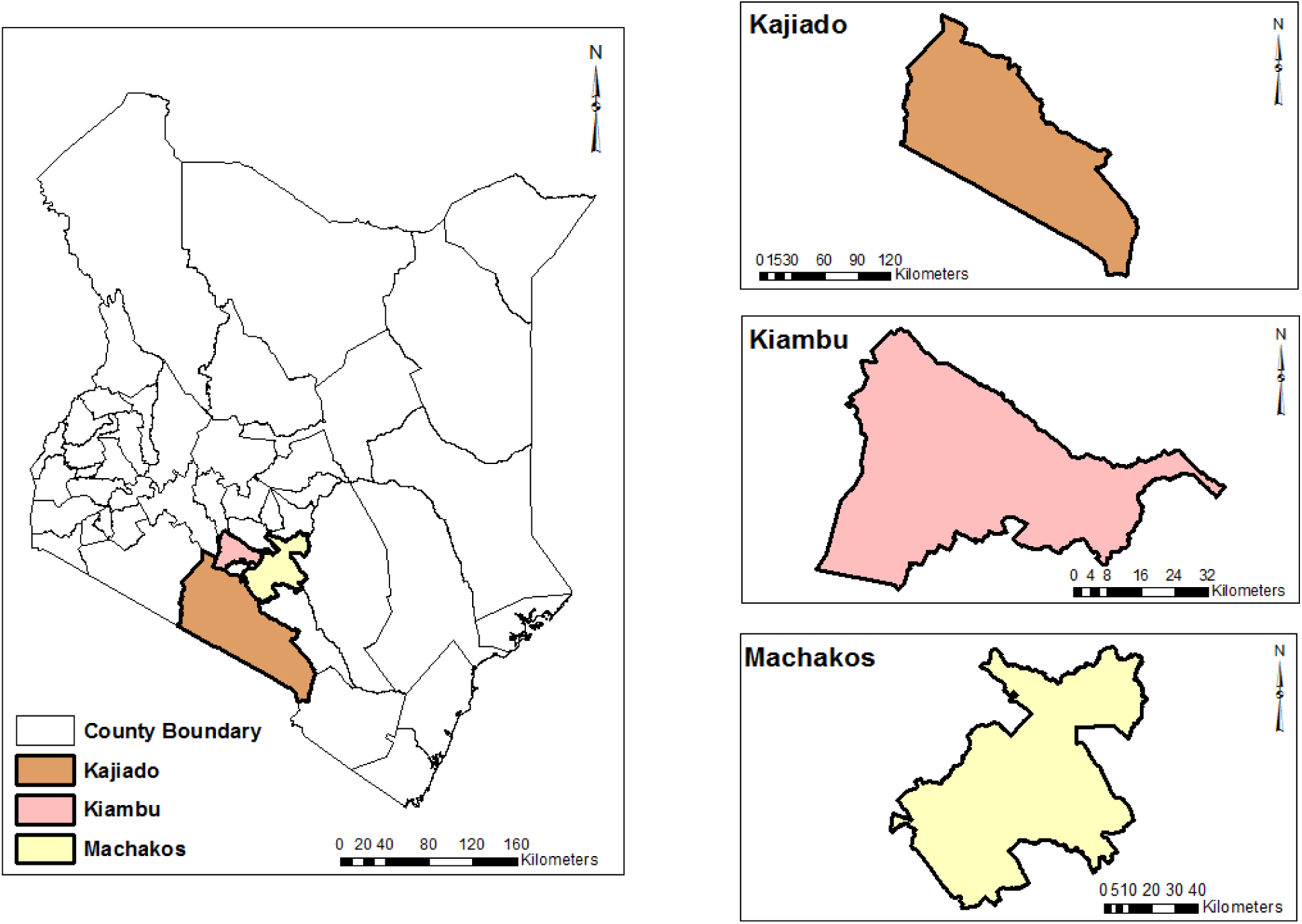
Map of Kenya showing the study site; Kajiado, Machakos and Kiambu Counties.

### Sample size determination, sampling design, and sample collection

This study employed a stratified random sampling technique, where the counties served as strata. In each stratum (Machakos, Kiambu and Kajiado), the purposive sampling technique was employed to select the 10 animal farms of interest, with the purpose being heavy or frequent utilization of antibiotics in the farms. A convenience sampling technique was employed in the tail end of the sample collection, where the samples were collected at the convenience of the farm owners. For downstream analysis, the samples were pooled into 18 groups where samples from the same source were pooled together.

According to anecdotal data collected from the three county Agriculture field extension officers, there are approximately between 100-150 farms in each of the three counties that rear cattle, pigs and poultry either in one farm or separately, making the total population about 330. The sample size was be calculated using the following formula by (Yamane, 1970).

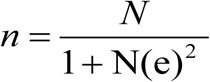

Where:

n =sample size

N is the population size 330

e= level of precision=0.05

therefore

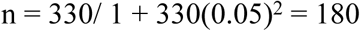

For each county, 10 manure samples from each of the three animals were sampled, along with the soil exposed to the manure, giving a total of 60 samples per county. In total, 180 samples were collected as per the calculated sample size.

### Ethical statement

Ethical approval and research permit was sought and granted from The National Commission for Science, Technology and Innovation (NACOSTI) in Kenya under License No: NACOSTI/P/23/29612.

### Extraction and Quantification of antibiotic residues in soil and manure samples

Extraction experiments were performed by weighing 1 g of each manure and soil sample into a sterile 50 mL polypropylene centrifuge tube, followed by the addition of 10 mL of extraction solvent. Each tube was vortexed vigorously using Thermo Scientific Digital Vortex Mixer-Vortexer, Cambridge Scientific, UK for one minute, followed by ultrasonic extraction using a sonotrode for 15 minutes. After sonication, the samples were centrifuged using Universal 320 Hettich centrifuge (Germany) at 3,000 × g for 5 minutes. A volume of 10 mL of the clear supernatant was carefully transferred into a 20 mL amber glass vial. The supernatant was then evaporated under a gentle stream of nitrogen gas at 40°C until the volume was reduced to approximately 5 mL. To further improve analyte retention during solid-phase extraction (SPE), samples were diluted with 13 mL of 0.1 M or 0.2 M EDTA solution.

Oasis HLB SPE cartridges were preconditioned sequentially with 5 mL of methanol and 5 mL of EDTA-McIlvaine buffer. The cartridges were loaded with the sample, then washed with 5 mL of bi-distilled water to remove residual impurities, and dried under a mild vacuum for 1 minute. Elution was performed in two steps using 2 × 3 mL of methanol, and the eluates were evaporated to dryness under nitrogen gas at 40°C, then reconstituted in 1 mL methanol The reconstituted samples were transferred into 2 mL Eppendorf tubes and centrifuged at 12,000 × g for 30 minutes. The clarified extracts were transferred into 1.5 mL amber vials and stored at -20°C until further analysis

Antibiotics were quantified using an LC-ESI-triple MS (API 3200, Applied Biosystems/MDS Sciex Instruments, Toronto, Canada) at lower and higher calibration range of 0-100 and 0-1000. The volume of methanol did not exceed 0.1% v/v of the total final volume of the stock solution prepared in 0.01 M CaCl_2_.

### Molecular detection and quantification of antibiotic-resistant genes

#### DNA extraction

DNA was extracted using the Quick-DNA™ Fecal/Soil Microbe Miniprep Kit from Zymo Research inc (Germany) following the manufactures instructions. Briefly, 150 mg of manure and 250 mg of soil sample was added to a ZR BashingBead™ Lysis Tube (0.1 & 0.5 mm). Subsequently, 750 µl of BashingBead™ Buffer was introduced into the tube. The tube was secured in a vortex mixer and vortexed for 40 minutes to lyse all the cells. Followed by centrifuged at ≥ 10,000 x g for 1 minute. 400 µl of the supernatant was transferred to a Zymo-Spin™ III-F Filter in a Collection Tube and centrifuged at 8,000 x g for 1 minute. Then, 1,200 µl of Genomic Lysis Buffer was added to the filtrate in the Collection Tube and mixed well. A total of 800 µl of this mixture was transferred to a Zymo-Spin™ IICR Column in a Collection Tube and centrifuged at 10,000 x g for 1 minute. The flow-through was discarded, and the step was repeated to ensure maximum binding. Next, 200 µl of DNA Pre-Wash Buffer was added to the Zymo-Spin™ IICR Column, which was placed in a new Collection Tube, and centrifuged at 10,000 x g for 1 minute. Following this, 500 µl of g-DNA Wash Buffer was added to the Zymo-Spin™ IICR Column and centrifuged at 10,000 x g for 1 minute. The Zymo-Spin™ IICR Column was then transferred to a clean 1.5 ml microcentrifuge tube, and 100 µl of DNA Elution Buffer was directly applied to the column matrix. The tube was centrifuged at 10,000 x g for 30 seconds to elute the DNA. A Zymo-Spin™ III-HRC Filter was placed in a clean Collection Tube, and 600 µl of Prep Solution was added. This assembly was centrifuged at 8,000 x g for 3 minutes. Finally, the eluted DNA was transferred to a prepared Zymo-Spin™ III-HRC Filter in a clean 1.5 ml microcentrifuge tube and centrifuged at exactly 16,000 x g for 3 minutes to complete the purification process.

#### Quantitative PCR for resistant genes

Absolute quantification of selected antibiotic resistance genes (ARGs)—*blaMOX, blaOXA, qnrB, aadA, dfrA1, sul1, tetQ, tetW,* and *ermB*—was performed using the qTOWER³ Auto real-time PCR system (Analytik Jena, Germany). Gene-specific primers targeting conserved regions of each ARG were designed (supplementary material 1) based on validated reference sequences from NCBI GenBank to ensure specificity and amplification efficiency. Primers were synthesized by General Biosystems Inc. (USA) using high-fidelity phosphoramidite chemistry and purified by high-performance liquid chromatography (HPLC). For absolute quantification, synthetic double-stranded DNA standards encompassing the full target amplicon regions were generated by GeneUniversal Inc. (USA). These standards were accurately quantified via spectrophotometry and serially diluted to construct standard curves for each gene. qPCR amplification reactions were carried out in 20 µL volumes using the innuMIX qPCR DSGreen Standard master mix (Analytik Jena, Germany), a dye-based mix optimized for high sensitivity and specificity. Each reaction consisted of: 10 µL of innuMIX qPCR DSGreen master mix, 1 µL of template DNA, 0.8 µL of forward primer, 0.8 µL of reverse primer, 7.6 µL of molecular-grade nuclease-free water. The PCR thermal cycling conditions were; initial denaturation at 95°C for 10 minutes, then 40 cycles of denaturation at 95°C for 20 seconds, annealing at 60°C for 1 minute and extension at 72°C for 30 seconds, with product melting between 60°C to 95°C for 15 minutes.

Following absolute quantification, relative quantification of the resistant genes was done by normalizing the ARGs copy numbers to the 16s rRNA gene copy number within the same sample. The normalization accounted for variation in microbial biomass and the total bacterial load which in turn allowed for an accurate cross sample comparison. This normalized data was subsequently used to evaluate the composition structure of the resistome, to estimate the ARG diversity and richness and to explore the relationship between ARGs and environmental factors. Multivariant analyses like non-metric multidimensional scaling (NMDS) and principal coordinate analysis (PCoA) were conducted to assess clustering by sample type or county.

#### Data Analysis

The average concentration of the antibiotic residues was calculated and visualized using bar charts to provide a summary of their distribution across samples. A three-way ANOVA was employed to analyse the variation in antibiotic concentrations across three categorical factors: sample type (manure and soil), animal species (cow, pig, and chicken), and county of origin (Machakos, Kiambu, and Kajiado). Absolute quantification of antimicrobial resistant genes (ARG) was performed, and the values were normalized to 16s rRNA gene copes to obtain relative quantification. Principal Coordinate Analysis (PCoA) was conducted to explore patterns in ARG composition, while spearman correlation analysis was employed to assess the co-occurrence and potential relationship among different ARGs.

## Results

### Occurrence of antibiotic residues in manure and soil

The total antibiotic concentration per county showed distinct spatial patterns in residue accumulation across Machakos, Kiambu, and Kajiado. Overall, Kiambu recorded the highest total antibiotic concentration, followed by Machakos, with Kajiado showing comparatively lower levels (supplementary material 2) (Figure 2). This suggests a heavier usage or persistence of veterinary antibiotics in Kiambu’s livestock production systems, potentially linked to more intensive pig and poultry farming activities.

**Figure 2:**
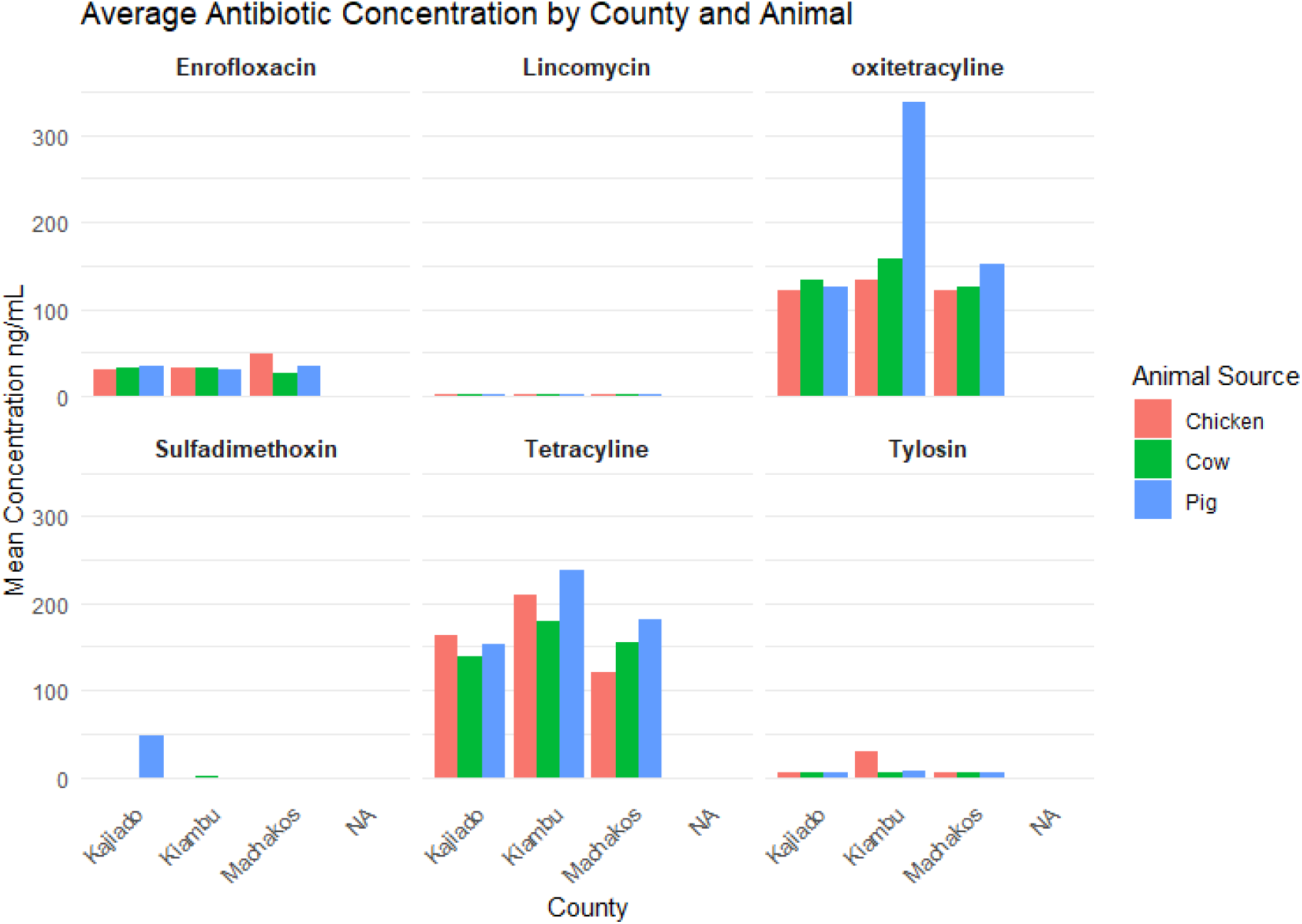
Average antibiotic concentration by counties (Kajiado, Kiambu and Machakos) and animal source (Cow, pig and Chicken)

Enrofloxacin was detected in all the counties, in both soil and manure samples originating from the three animals manure showing a higher concentration than soils. Chicken manure from Machakos exhibited the highest median concentration (∼65 ng/mL) and soil laced with cow manure from Machakos showed the lowest concentration, approaching zero. Lincomycin concentration was detected in low quantities across the counties and in all the animals with a concentration ranging from 0 to about 3.41 ng/mL from soil laced with cow manure in Kajiado County. Oxytetracycline showed a high concentration in all the tested samples across the three counties. The highest average concentration was recorded at over 665 ng/mL in pig manure from Kiambu County. Tetracycline also showed consistent high concentration across all samples pig manure from Kiambu County showing the highest concentration over 200 ng/mL. As shown in figure 2 above, sulfadimethoxin was below detectable limits in most samples except manure from pigs in Kajiado County whose concentration averaged at 127 ng/mL. Tylosin showed low concentration across the samples with the highest concentration being chicken samples from Kiambu at an average of 45 ng/mL.

**Figure 3:**
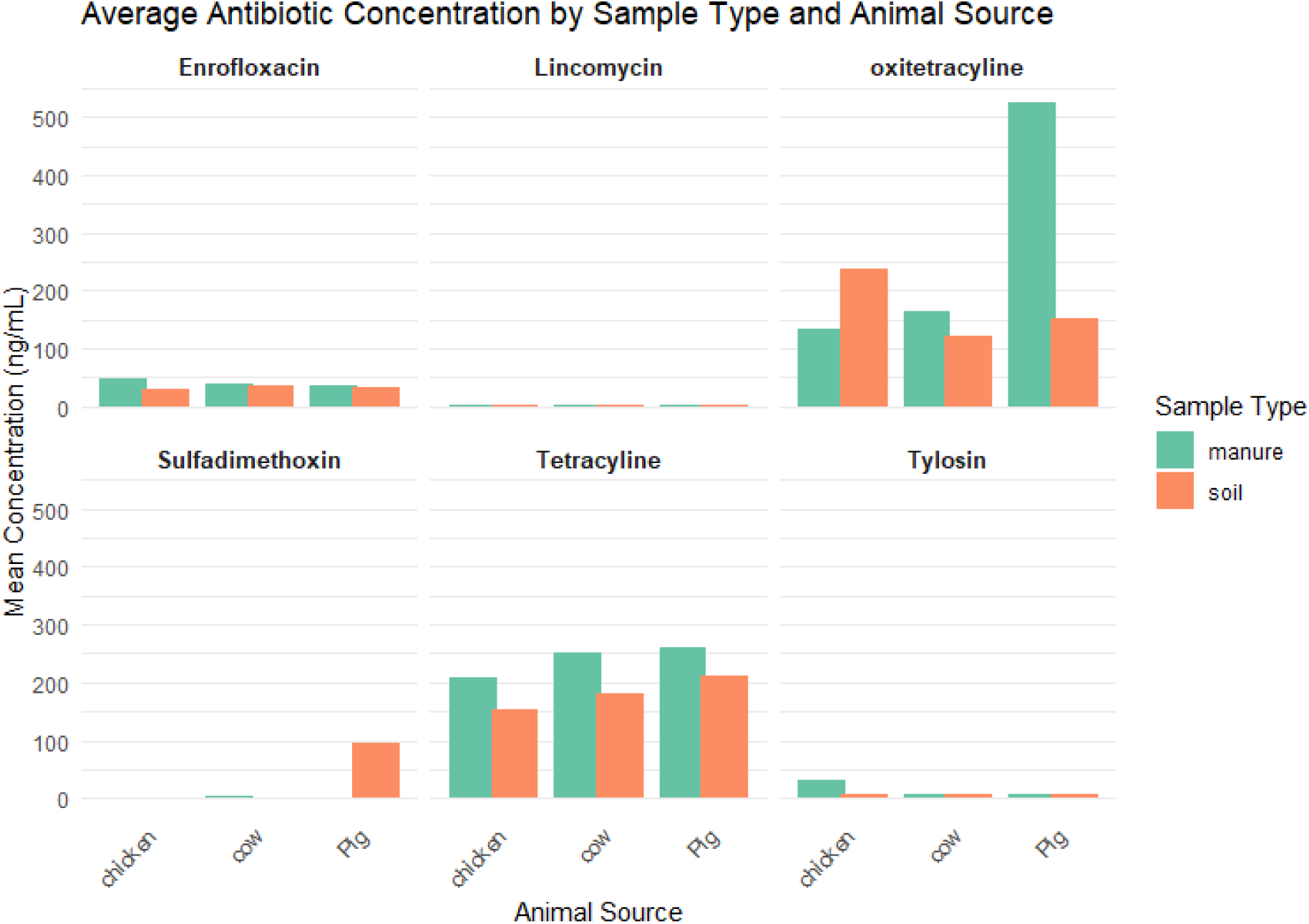
Average concentration of antibiotics by sample type (manure and soil) and animal source (cow, pig and chicken)

Three-way ANOVA results generally showed that the differences in antibiotic concentrations from the animal source (cow, pig and chicken) are not a significant factor in the antibiotic concentration. Sulfadimethoxin and oxytetracycline showed marginal significance with a P-value of 0.0904 and 0.1003, respectively. Sample type (manure and soil) did not show a strong effect on antibiotic concentration only tetracycline showed a marginal significance with a p-value of 0.0785, showing a possible influence of sample type on its residue levels. When the county (Machakos, Kiambu and Kajiado) was analyzed, it was shown that only oxytetracycline (P = 0.0421) was significant, which showed differences in residues across counties. Two-way variations were considered and animal source and sample type showed no significant variation for any antibiotics only oxytetracycline approached significance (p = 0.0901). Animal source and county only showed statistical significance for sulfadimethoxin (P = 0.044) and sample type and county did not show significant effects. Three-way interactions (animal source, sample type and county) were significant for oxytetracycline (P = 0.0453) and approaching significance for sulfadimethoxin (P = 0.0453) as shown in Table 1 below.

**Table 1:** Summary of three-way ANOVA results evaluating the effects of animal source, sample type and county on antibiotic concentration across seven antibiotics residues.

| Source of Variation | Antibiotic | Df | Sum Sq | Mean Sq | F value | Pr(>F) |
| --- | --- | --- | --- | --- | --- | --- |
| animal_source | Amoxicillin | 2 | 143 | 71.4 | 0.239 | 0.788 |
|  | Enrofloxacin | 2 | 143 | 71.4 | 0.239 | 0.788 |
|  | Lincomycin | 2 | 0.42 | 0.2122 | 0.086 | 0.917 |
|  | Tylosin | 2 | 149 | 74.75 | 0.836 | 0.439 |
|  | Sulfadimethoxin | 2 | 3990 | 1995 | 2.513 | 0.0904 |
|  | Oxytetracycline | 2 | 78,984 | 39,492 | 2.401 | 0.1003 |
|  | Tetracycline | 2 | 17,280 | 8,640 | 0.853 | 0.4319 |
| sample_type | Amoxicillin | 1 | 488 | 488.3 | 1.635 | 0.206 |
|  | Enrofloxacin | 1 | 488 | 488.3 | 1.635 | 0.206 |
|  | Lincomycin | 1 | 0.05 | 0.0512 | 0.021 | 0.886 |
|  | Tylosin | 1 | 169 | 169 | 1.889 | 0.175 |
|  | Sulfadimethoxin | 1 | 1875 | 1875.2 | 2.362 | 0.1301 |
|  | Oxytetracycline | 1 | 26,508 | 26,508 | 1.611 | 0.2097 |
|  | Tetracycline | 1 | 32,585 | 32,585 | 3.216 | 0.0785 |
| county | Amoxicillin | 2 | 17 | 8.5 | 0.029 | 0.972 |
|  | Enrofloxacin | 2 | 17 | 8.5 | 0.029 | 0.972 |
|  | Lincomycin | 2 | 0.24 | 0.1218 | 0.05 | 0.952 |
|  | Tylosin | 2 | 308 | 154.03 | 1.722 | 0.188 |
|  | Sulfadimethoxin | 2 | 3990 | 1995 | 2.513 | 0.0904 |
|  | Oxytetracycline | 2 | 110,599 | 55,300 | 3.362 | 0.0421 |
|  | Tetracycline | 2 | 42,651 | 21,326 | 2.105 | 0.1318 |
| **animal_source × sample_type** | Amoxicillin | 2 | 51 | 25.7 | 0.086 | 0.918 |
|  | Enrofloxacin | 2 | 51 | 25.7 | 0.086 | 0.918 |
|  | Lincomycin | 2 | 0.02 | 0.0102 | 0.004 | 0.996 |
|  | Tylosin | 2 | 317 | 158.57 | 1.773 | 0.18 |
|  | Sulfadimethoxin | 2 | 4123 | 2061.6 | 2.597 | 0.0838 |
|  | Oxytetracycline | 2 | 82,835 | 41,417 | 2.518 | 0.0901 |
|  | Tetracycline | 2 | 9,713 | 4,857 | 0.479 | 0.6218 |
| **animal_source × county** | Amoxicillin | 4 | 993 | 248.1 | 0.831 | 0.511 |
|  | Enrofloxacin | 4 | 993 | 248.1 | 0.831 | 0.511 |
|  | Lincomycin | 4 | 0.13 | 0.033 | 0.013 | 1 |
|  | Tylosin | 4 | 363 | 90.76 | 1.015 | 0.408 |
|  | Sulfadimethoxin | 4 | 8358 | 2089.6 | 2.632 | 0.044 |
|  | Oxytetracycline | 4 | 128,912 | 32,228 | 1.959 | 0.1139 |
|  | Tetracycline | 4 | 14,448 | 3,612 | 0.357 | 0.8384 |
| **sample_type × county** | Amoxicillin | 2 | 675 | 337.7 | 1.131 | 0.33 |
|  | Enrofloxacin | 2 | 675 | 337.7 | 1.131 | 0.33 |
|  | Lincomycin | 2 | 0.07 | 0.0358 | 0.015 | 0.986 |
|  | Tylosin | 2 | 333 | 166.68 | 1.864 | 0.165 |
|  | Sulfadimethoxin | 2 | 4123 | 2061.6 | 2.597 | 0.0838 |
|  | Oxytetracycline | 2 | 23,719 | 11,859 | 0.721 | 0.4909 |
|  | Tetracycline | 2 | 9,669 | 4,835 | 0.477 | 0.6231 |
| **animal_source × sample_type × county** | Amoxicillin | 4 | 320 | 79.9 | 0.268 | 0.898 |
|  | Enrofloxacin | 4 | 320 | 79.9 | 0.268 | 0.898 |
|  | Lincomycin | 4 | 0.77 | 0.1919 | 0.078 | 0.989 |
|  | Tylosin | 4 | 644 | 161.05 | 1.801 | 0.142 |
|  | Sulfadimethoxin | 4 | 7874 | 1968.4 | 2.48 | 0.0547 |
|  | Oxytetracycline | 4 | 171,891 | 42,973 | 2.612 | 0.0453 |
|  | Tetracycline | 4 | 19,959 | 4,990 | 0.492 | 0.7412 |

In terms of specific antibiotic residues, amoxicillin, enrofloxacin and lincomycin showed no statistically significant effects observed from any factor or interaction. Tylosin also showed no significant predictors but multiple factors had moderately high F-values which suggests possible trends. Sulfadimethoxin exhibited a near-significance interaction consistently, particularly between counties. Oxytetracycline displayed main and interaction effects across various factors, whereas whole tetracycline only showed marginal significance with sample type as shown in Table 1 below.

### Quantification of Antimicrobial Resistance Genes

In Kajiado County, manure samples showed a substantially higher ARG abundance as compared to soil samples. Conversely, Kiambu and Machakos exhibited high levels ARG in soil samples. (figure 4)

**Figure 4:**
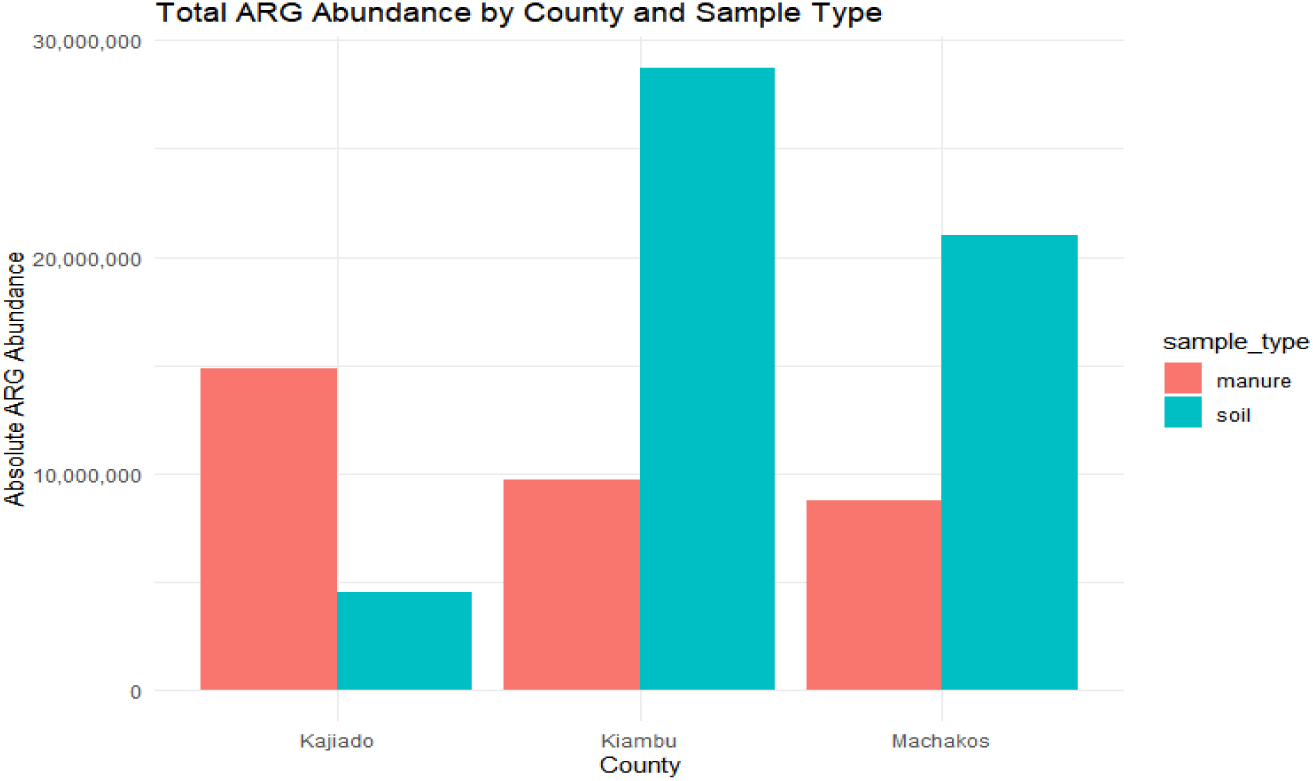
Total ARG abundance by county and sample type

In terms of specific genes, the *aadA* gene showed the highest abundance among all the ARGs analysed particularly in Machakos which suggest possible high usage of aminoglycosides in the area. *ermB* gene was also present in high values in the same county. *Sul1* which is associated with sulphonamide resistance, showed a considerable variation (supplementary material 3).

On the other hand, *blaMOX* and *QnrB* displayed a very low abundance across all counties,. *tetW* and *tetQ* genes, which are both tetracycline resistance genes showed moderate levels with slightly higher distribution in Machakos. As shown in Figure 5 above, which indicates a distinct spatial heterogeneity in ARG distribution with Machakos showing generally higher levels of several key resistant genes like *aadA, sul1*and *blaOXA*.

**Figure 5:**
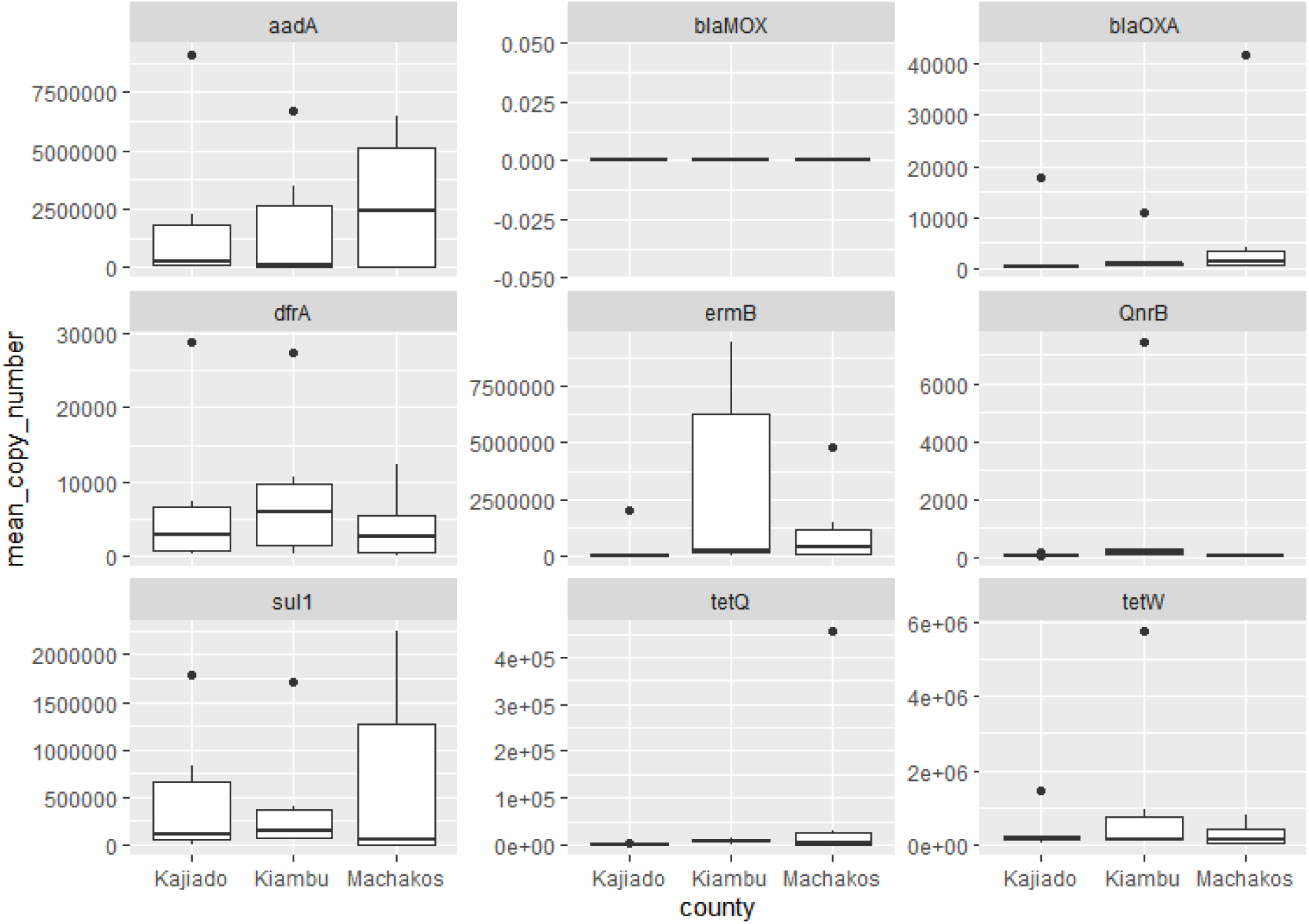
Distribution of mean copy numbers for individual antibiotic resistance genes (ARG) across the three counties

### Relative quantification

Relative quantification was conducted by dividing the absolute abundance of each ARG by the abundance of the 16s rRNA gene in the same sample. This ratio gave the relative abundance of the ARGs per bacterial cell which provided a standardized measure of gene prevalence withing each of the samples tested. The findings of this study showed that in Kajiado County, *aadA*, *ermB*, and *tetW* genes exhibited the highest abundance levels while *blaOXA and sul1* were relatively low. In Kiambu County, there was a general reduction in abundance with a measurable presence of *ermB* and *QnrB.* Machakos County also showed similar patterns but with *aadA* and *tetW* showing higher levels as shown in figure 6 above.

**Figure 6:**
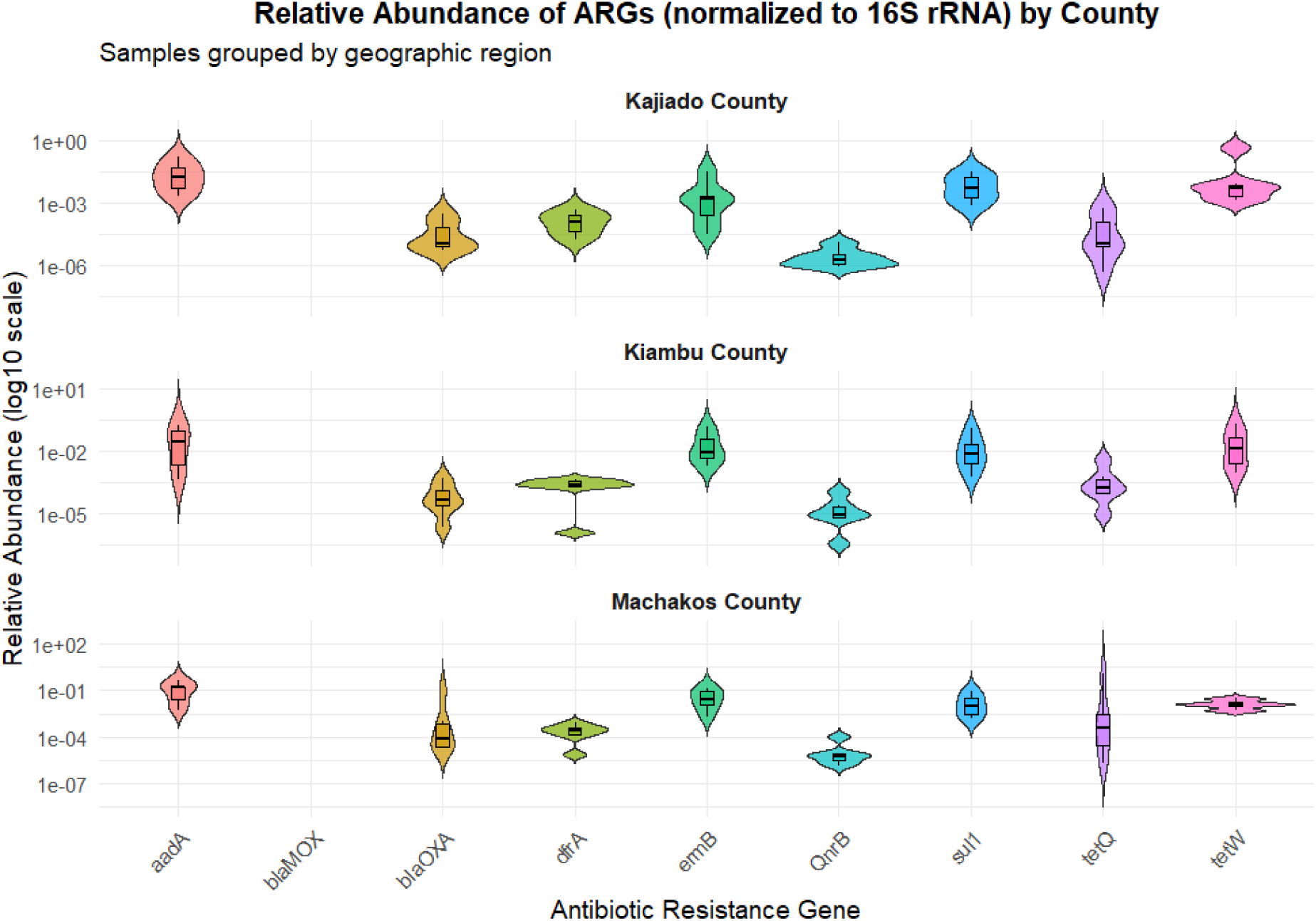
Relative abundance of antibiotic resistance genes (ARGs) across individual samples, normalized to 16s rRNA gene copy numbers (ARG/16s)

The relative abundance of the ARGs were analyzed using the sample type (manure or soil) and the animal type (cow, pig and chicken) in the cow samples the *aadA, ermB, sul1* and *tetQ* showed relative high abundance in manure compared to soil samples. The soil samples retained detectable levels of *ermB* and *tetQ* and the *blaMOX* and *blaOXA* exhibited lower abundance. In the pig samples, *ermB, tetQ* and *tetW* were higher particularly in manure compared to soil samples which showed a reduced ARG abundance, but *ermB* and *qnrB* remained consistently high presence while *blaOXA* and *sul1* displayed low to moderate abundance across both samples. In chicken samples, *aadA, ermB* and *tetQ* were dominant across both manure and soil samples with a higher variability in manure. *blaOXA* and *qnrB* showed low abundance across the samples as shown in figure 7 above.

**Figure 7:**
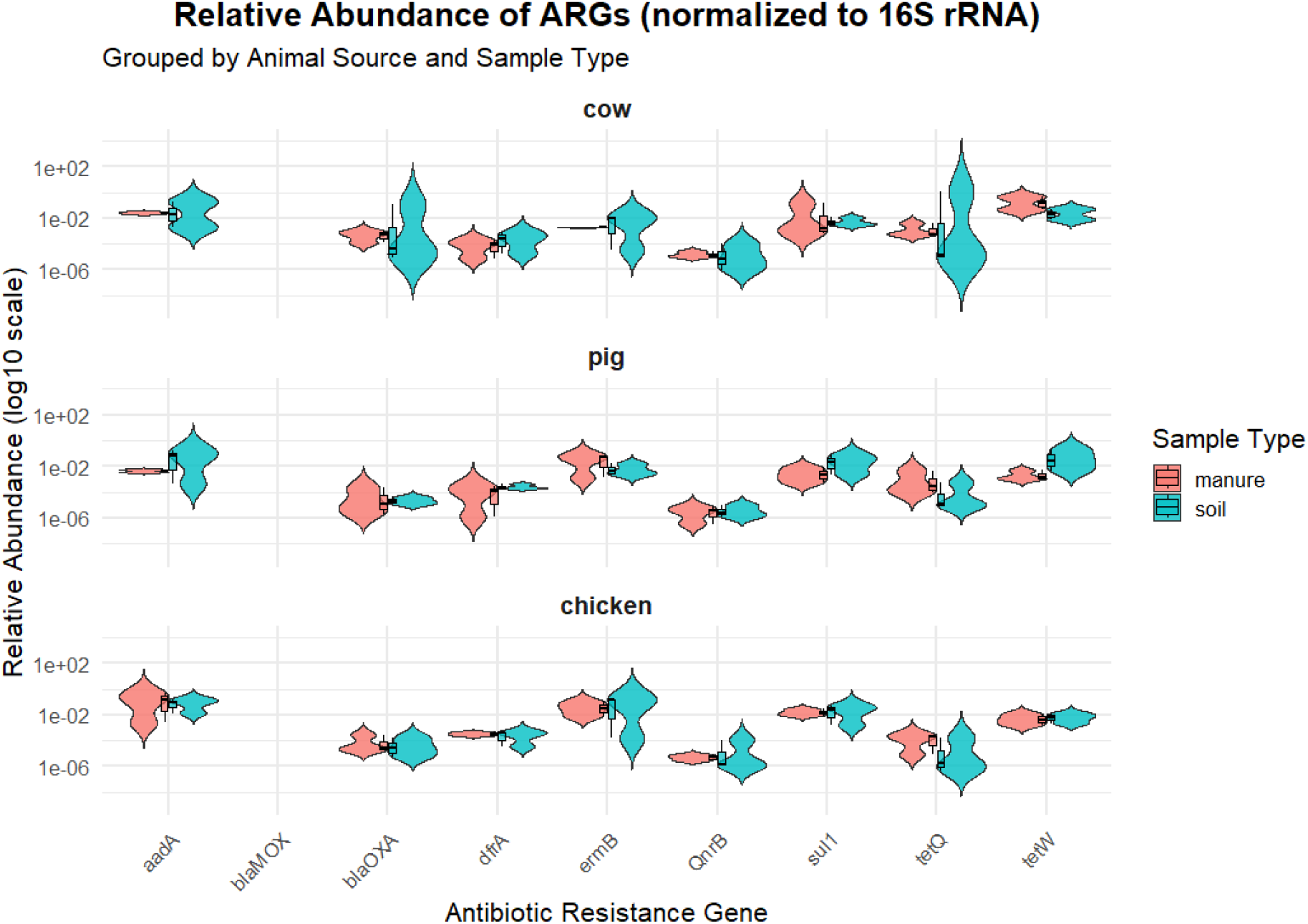
Relative abundance of ARGs normalized to 16s rRNA gene and grouped by the animal’s source of the samples and sample type

### Principal Component Analysis (PCA)

Principal coordinate analysis showed that most samples share similar ARG profiles, but a few samples were clearly distinct due to the environmental and biological factors. As shown in figure 8 below most samples were tightly clustered near the point of origin (PC1 ≈ 0, PC2 ≈ 0) which suggest a similar composition among those samples.

**Figure 8:**
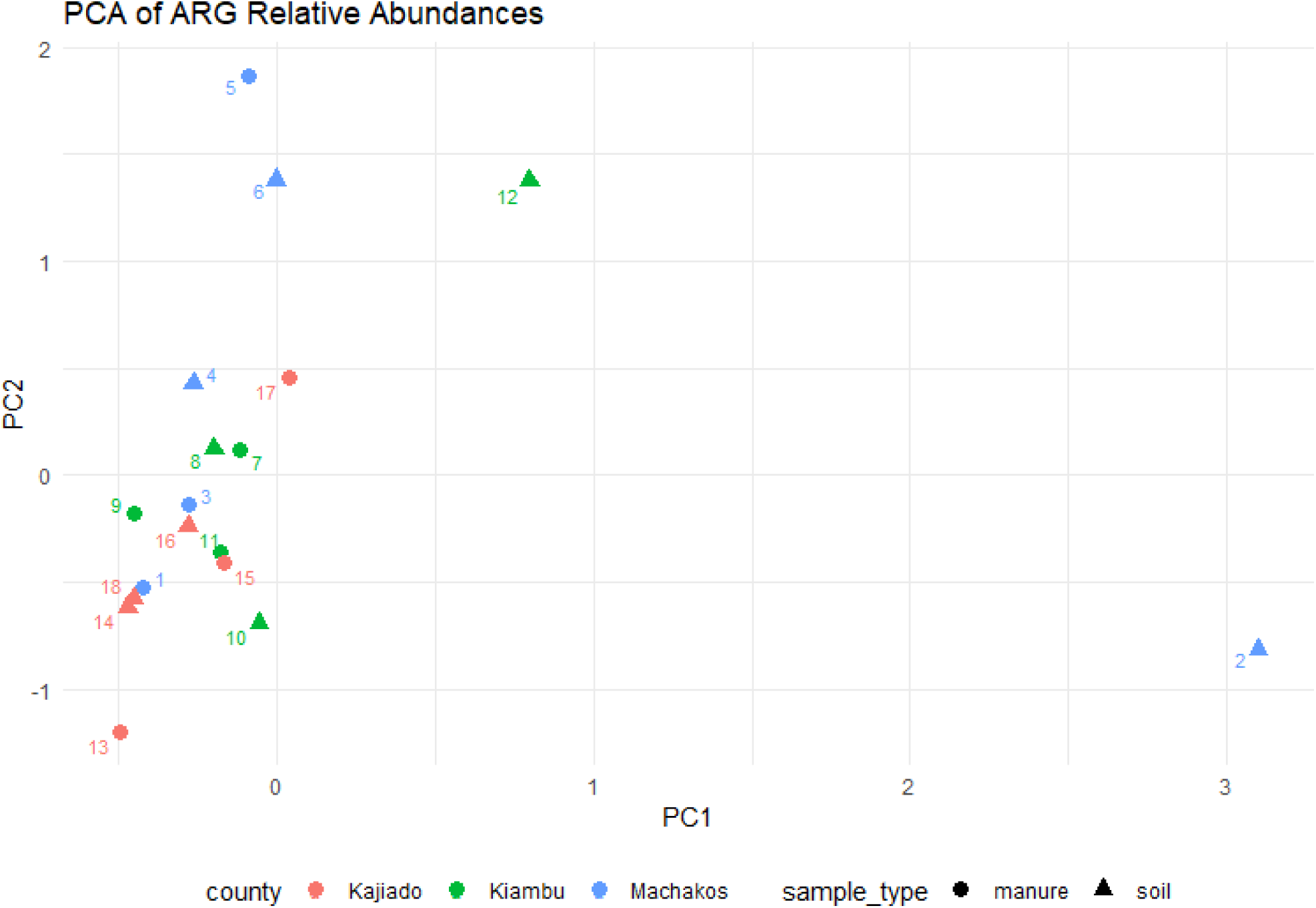
The principal component analysis (PCA) plot illustrates the variation of relative abundance of antibiotic resistance genes (ARGs) across various samples types (manure and soil) from three counties (Kajiado, Kiambu and Machakos).

The clustering patterns indicate that majority of the samples, especially those from Kajiado(red) are tightly grouped near the origin, which indicates similar ARG profiles and indicates minimal variability in resistance gene abundance within the county. In contrast, samples from Machakos (blue) and Kiambu (green) exhibit greater dispersion across both axes, with several soil samples showing clear outliers (samples 2,5,6, and 12), which highlights potential unique or elevated ARG profiles in these samples. The soil samples show a wider separation in PCA space than manure, which suggests that soil harbors a more heterogeneous resistome.

**Figure 9:**
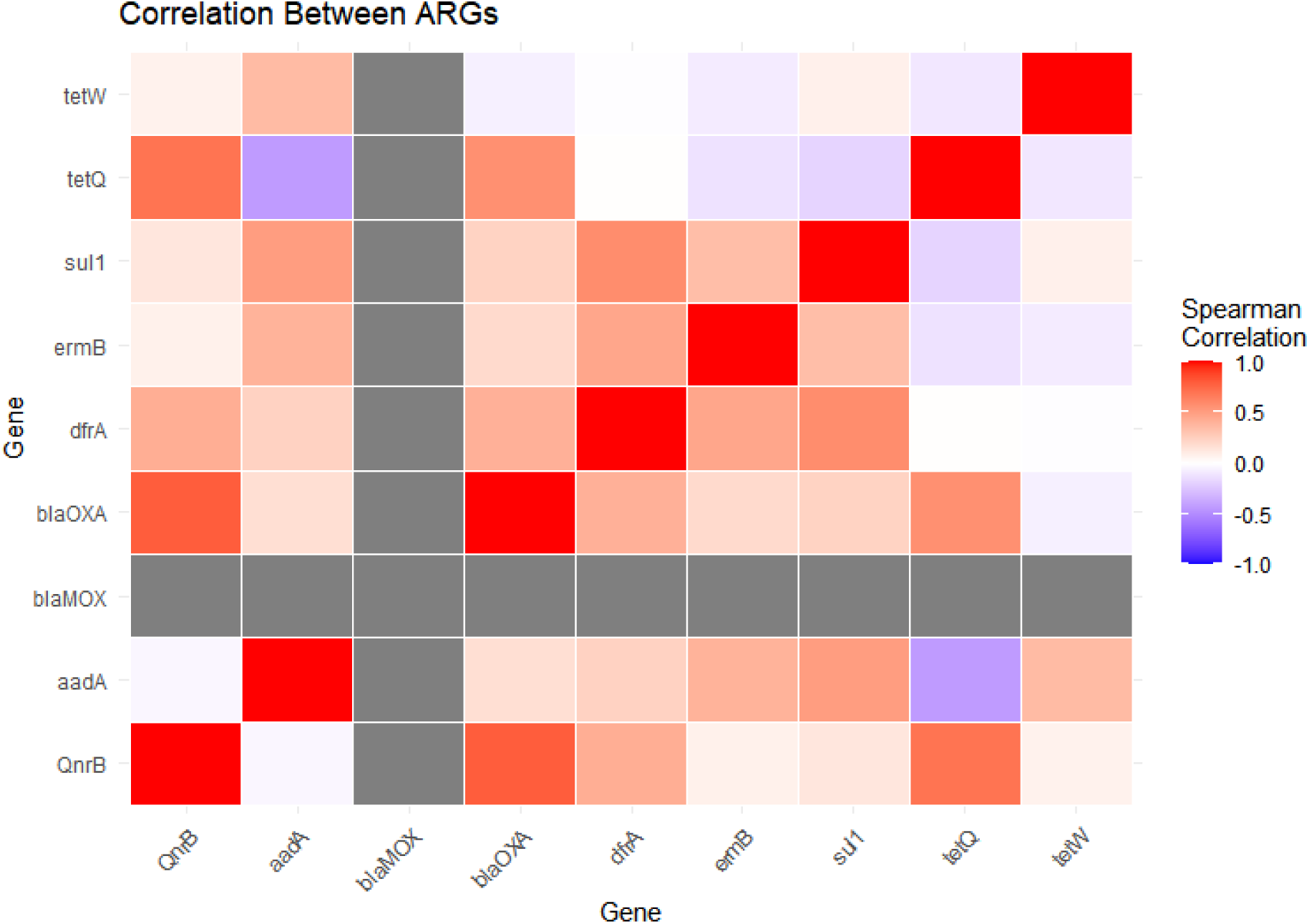
Spearman correlation coefficient between abundance of various antibiotic-resistant genes across multiple samples

Spearman correlation was conducted to illustrate the coefficients between the relative abundance of the ARGs across the samples. Several AGs exhibited strong positive correlation, notably *ermB* and *sul1*, *aadA* and *QnrB* and *tetQ* and *tetW,* which suggested that these genes may co-occur within the same microbial communities. *TetQ* and *tetW* positive correlation for example, may be due to exposure to tetracycline antibiotics in the same environment. There are a few negative correlations as shown in the heatmap in figure 10 above such as between *tetQ* and *aadA*, which indicates differential distributions or regulatory patterns.

## Discussion

The spatial analysis of antibiotic residues revealed a significant disparity among the counties, with Kiambu County recording the highest levels of antibiotic concentrations, followed by Machakos and Kajiado. These findings are supported by a previous study conducted in Kiambu among poultry farmers that showed a widespread, over-the-counter purchase and use of antibiotics (Kariuki et al., 2023). Tetracyclines were observed to be the most frequently occurring antibiotic, followed by enrofloxacin and tylosin. This is in agreement with a previous report on antibiotic use by poultry farmers in Kiambu county, which stated that tetracycline usage (34%) was higher than sulphonamides and Tylosin (10-15%) (Kariuki et al., 2023). Among the antibiotics tested, tetracyclines (especially oxytetracycline) were the most dominant across all samples, with concentrations above 1000 ng/mL in pig manure from Kiambu. Oxytetracycline has been heavily used in animal husbandry, where over 81% of cattle farmers reported its use (Kisoo et al., 2023). Tetracyclines have been identified as environmental pollutants worldwide (Amangelsin et al., 2023), and residues of tetracycline antibiotics have been found in agricultural soils and vegetables at levels as high as 171.07–660.20 μg kg−1 for tetracycline, 25.38–345.78 μg kg−1 for oxytetracycline, and 170.77–707.47 μg kg−1 for chlortetracycline (Xu et al., 2023). In contrast, our findings showed low or undetectable levels of sulphonamides and macrolides, potentially due to more rapid degradation of sulphonamides compared to others (Paumelle et al., 2021). The relatively high occurrence of tetracycline residues and prevalence of the associated *tetQ* and *tetW* ARGs in both manure and soil samples is partly because tetracyclines are difficult to metabolize in human and animal digestive systems. Between 50-80% of the molecules are excreted into the environment, where they persist (Fiaz et al., 2021). Manure samples, especially from pig, consistently showed high residue levels, emphasizing the risk posed by untreated or poorly managed animal waste. Interestingly, soil samples in several cases displayed comparable or even higher concentrations relative to their manure counterparts, suggesting residue accumulation over time and potential environmental amplification. These findings emphasize the environmental persistence of certain antibiotics and the need to assess their fate in soil, especially when organic fertilizers are used repeatedly.

The absolute and relative quantification of the ARGs revealed a widespread dissemination across the counties, with very clear antibiotic use and environmental persistence. Kiambu soil samples exhibited the highest ARG leads of up to 30 million copies, whereas Kajiado manure samples contained markedly more ARGs than the corresponding soils, which shows county-specific dynamics in ARG mobility and environmental loading. Among the ARGs analysed, *aadA* (aminoglycoside resistance), *ermB* (macrolide resistance), and *sul1* (sulphonamide resistance) showed the highest absolute abundances, particularly in Machakos and Kiambu. This distribution reflects both the prevalence and selective pressure of the corresponding antibiotic in these regions. Conversely, *blaMOX* (beta-lactam resistance) and *qnrB* (quinoline resistance) were not detected rarely detected, respectively, indicating either limited exposure to these antibiotics’ classes or poor environmental persistence of their genetic determinants. The detection of *tetQ* and *tetW* genes that confer resistance of tetracycline further mirrored the chemical residue data. These genes were particularly prevalent in samples from Machakos, underscoring the consistent relationship between tetracycline residue load associated resistome.

Normalization of ARG abundance to *16s rRNA* gene copy number provided a clearer view of ARG prevalence independent of the microbial biomass. Strikingly, *tetQ* approached a 1:1 ratio with *16s rRNA* in some samples. This is particularly alarming and may be indicative of intense selective pressure in certain microenvironments. The other genes *aadA, ermB* and *dfrA* showed moderate to high relative abundance, which suggests a widespread dissemination across the microbial communities (Reference). Hierarchical clustering and heatmap analysis confirmed that *tetQ* and *tetW* were the most prevalent and co-occurring genes across samples. Principal coordinate analysis (PCoA) further demonstrated clustering of most samples, indicating similarity in ARG profiles, although a few samples exhibited distinct resistome composition.

Spearman correlation analysis revealed several strong positive relationships, including between *ermB* and *sul1*, *aadA* and *qnr*, and *tetQ* and *tetW.* These findings suggest that co-selection and horizontal gene transfer maybe shaping ARG co-occurrence patterns. Similarly, negative correlations, such as those between *tetQ* and *aadA*, may reflect ecological trade-offs or differing niches among the resistant microbial communities.

## Conclusion

This study demonstrates a clear spatial heterogeneity in antibiotic residues and antibiotic residue distribution and antibiotic resistance genes (ARG) prevalence across Machakos, Kiambu and Kajiado Counties, revealing a significant link between antibiotic use practices and environmental contamination. Kiambu County showed the highest antibiotic residues residue loads, especially tetracyclines like oxytetracycline, corroborating prior evidence of intensive and mostly unregulated use inn livestock farming. The high concentration observed in manure, particularly from pigs, and the comparable elevated levels in soils indicate that antibiotics residues persist and accumulate in the environment, likely due to continuous manure application and to the poor degradation characteristic of tetracyclines. The persistence contributes to sustained selective pressure on environmental microbial community, thereby promoting the maintenance and proliferation of ARGs.

The Resistome analysis further substantiates this observation, showing a widespread presence of ARGs across all the counties with notable county-specific variation. The high abundance of *aadA, ermB sul1 tetQ* and *tetW* genes reflect the extensive use and persistence of aminoglycoside, macrolides, sulphonamide and tetracycline antibiotics. The strong alignment between the residue data and the prevalence of associated resistance genes selection highlights the ecological linkage between antibiotic contamination and genetic resistance selection. Particularly alarming is the near 1:1 ratio of *tetQ* to 16s rRNA gene copies in certain samples, suggesting pervasive tetracycline resistance and intense selective pressure within microbial populations. The positive correlation among multiple ARGs further indicate that co-selection and horizontal gene transfer are actively shaping ARG co-occurrence and dissemination patterns. Collectively these findings underscore the role of animal production systems as a significant reservoirs and amplifiers of antibiotic resistance within the environment.

Given the findings of this study, we recommend strengthening the regulation and stewardship of antibiotics in livestock production, including a strict prescription only antibiotic sales and stringent monitoring of antibiotics usage in animal husbandry. The animal manure needs more sustainable treatment and management such as composting, anaerobic digestion and bioremediation before using in crop production. These processes can minimize antibiotic residues and ARG loads which in tun will minimize their transfer into the agricultural soils. There is need for the government through the ministry of agriculture and the ministry of environment to establish a systematic national surveillance framework from tracking antibiotic residues and ARGs across environmental matrices like manure, soil water etc. Such a system should include molecular, chemical and geospatial data to identify high-risk zones and inform policy decisions. There is need for integrated efforts using the one health approach that brings environmental, agriculture and health policies in order to mitigate the environmental degradation due to misuse of antibiotics.

## Acknowledgement

The authors would like to acknowledge the Alexander von Humbold Foundation for funding this work. Additionally, we would like to acknowledge and thank the technical research assistants who helped in the research work. These include Njenga and Grace both from the Physical Sciences Department at Machakos University wo helped at the sampling stage of the study and Elvira Sieberger and Petra Ziegler from the Soil Science Department at Trier University wo helped with the laboratory work.

